# Dual control of meiotic crossover patterning

**DOI:** 10.1101/2022.05.11.491364

**Authors:** Stéphanie Durand, Qichao Lian, Juli Jing, Marcel Ernst, Mathilde Grelon, David Zwicker, Raphael Mercier

**Affiliations:** Department of Chromosome Biology, Max Planck Institute for Plant Breeding Research, Carl-von-Linné-Weg 10, 50829 Cologne, Germany; Max Planck Institute for Dynamics and Self-Organization, Am Faßberg 17, 37077 Göttingen, Germany; Université Paris-Saclay, INRAE, AgroParisTech, Institut Jean-Pierre Bourgin (IJPB), 78000, Versailles, France

## Abstract

Most meiotic crossovers (COs), called class I crossovers, are produced by a conserved pathway catalyzed by the ZMM proteins; COs are limited in number, typically to 1–3 per chromosome, and are prevented from occurring close to one other by crossover interference^1-3^. In many species, CO number is subject to dimorphism between males and females, and a lower CO number is associated with shorter chromosome axes and stronger interference^4^. How the patterning of COs is imposed, however, remains poorly understood. Here, we show that overexpression of the ZMM protein HEI10 increases COs and reduces crossover interference but maintains sexual dimorphism; shorter axes length in female meiosis is still associated with fewer COs and stronger interference than in male meiocytes. Disrupting the synaptonemal complex (SC) by mutating *ZYP1* also leads to an increase in class I COs but, in contrast, abolishes interference and disrupts the link between chromosome axis length and COs, with female and male meiocytes having the same CO frequency despite different axis lengths. Combining HEI10 overexpression and *zyp1* mutation leads to a massive increase in class I COs and absence of interference, while axes lengths are still unaffected. These observations support, and can be effectively predicted by, a recently proposed coarsening model^5,6^ in which HEI10 diffusion is funneled by the central element of the SC before coarsening into large, well-spaced CO-promoting droplets. Given the conservation of the components, this model may account for CO patterning in many eukaryotes.

## Introduction

A hallmark of sexual reproduction is the shuffling of homologous chromosomes by meiotic crossovers (COs). COs are produced by the repair of DNA double-strand breaks through two biochemical pathways: Class I COs are produced by a meiotic-specific pathway catalyzed by the ZMM proteins (*Saccharomyces cerevisiae* Zip1-4, Msh4-5, and Mer3) and represent most COs; Class II COs originate from a minor pathway that uses structure-specific DNA nucleases also implicated in DNA repair in somatic cells. Despite a vast excess of initial double-strand breaks, the number of resulting COs is limited, typically to one to three per chromosome pair. Class I COs are subject to additional tight constraints: At least one class I CO occurs per chromosome pair at each meiosis, the so-called obligate CO that ensures balanced chromosome distribution. Class I COs are also prevented from occurring next to each other on the same chromosome, a phenomenon called crossover interference. How this interference is achieved mechanistically has been debated for over a century^1,3,7-9^.

One specific unresolved question is the role of the synaptonemal complex (SC) in crossover interference. The SC is a zipper-like tripartite structure composed of two lateral chromosome axes, along which arrays of chromatin loops of each of the two homologous chromosomes are anchored, and a central part consisting of transverse filaments that connect the axes all along their length at meiotic prophase. Assessing the role of the SC in interference is difficult, because in many organisms the transverse filament protein is essential for the formation of class I COs^2^. One notable exception is *Arabidopsis thaliana*, where the transverse filament protein is not required for class I CO formation, providing a unique opportunity to analyze the role of the SC in CO patterning. In the *zyp1* mutant, class I COs form at a higher frequency than wild type and completely lack interference, demonstrating that the central element of the SC is, directly or indirectly, essential for imposing CO interference in Arabidopsis^10,11^. Reduced expression of the transverse element in *C. elegans* and specific mutations of the SC component that uncouple SC and CO formation in budding yeast lead to a reduction of interference, supporting a conserved role of the SC in imposing interference^12-15^. Interestingly in some species, such as humans or Arabidopsis, CO number differs in males and females. This heterochiasmy correlates with axis/SC length, with the number of COs proportional to axis length^4,16,17^. CO interference appears to propagate at a similar axis/SC distance (µm) in both sexes, which means that interference acts over greater genomic ranges (DNA) in the sex with a shorter axis/SC^17,18^, an observation which shows that the relevant space for the mechanism of interference is the axis/SC length.

A model was recently elaborated to account for class I CO patterning and interference, based on diffusion of the ZMM protein HEI10 (ZHP-3/4 in *C*.*elegans*) within the SC and a coarsening process leading to well-spaced CO-promoting HEI10 droplets^5,6^. HEI10, which encodes an E3 ubiquitin ligase, initially forms multiple small foci along the SC and is progressively consolidated into a small number of large condensates that co-localize with CO sites in diverse species^19-22^. Further, as predicted by the model, CO numbers depend on HEI10 dosage in Arabidopsis^6,23^. Interference is abolished in the absence of the transverse element of the synaptonemal complex ZYP1^10,11^, which is compatible with the idea that diffusion of HEI10 along the central part of the SC underlies crossover patterning and interference.

Here, we explored the mechanisms of CO patterning in Arabidopsis by analyzing the combinatory effects of axis/SC length (male vs. female), modification of HEI10 dosage, and disruption of the SC on COs. We notably show that overexpressing HEI10 in *zyp1* completely deregulates class I COs, with a massive increase of their number in both females and males. Our results support the model in which HEI10 coarsening by diffusion along the central element of the SC mediates CO patterning and imposes CO interference.

## Results and discussion

To decipher CO control, we studied the number and distribution of COs in both female and male meiosis when overexpressing HEI10 (well-characterized C2 line^23^), in the absence of the synaptonemal complex (*zyp1*), and in combination. We measured the number of class I COs in meiocytes by counting the number of MLH1-HEI10 co-foci at diplotene. In the pure line Col we analyzed six genotypes: wild type and *zyp1-1* combined with three dosages or HEI10 (wild type, heterozygous or homozygous for the HEI10^oe^ C2 transgene). In the Col/L*er* F1 hybrid, we analyzed four genotypes: wild type and *zyp1-1/zyp1-6* combined with two dosages of HEI10 (wild type and heterozygous for the C2 HEI10 transgene) (Figure 1A-B). The four hybrid genotypes were also used to characterize the crossover number and distribution by sequencing populations derived from female and male crosses to Col (Figure 1C–F, Figure 2, Figure S1–4).

**Figure 1.**
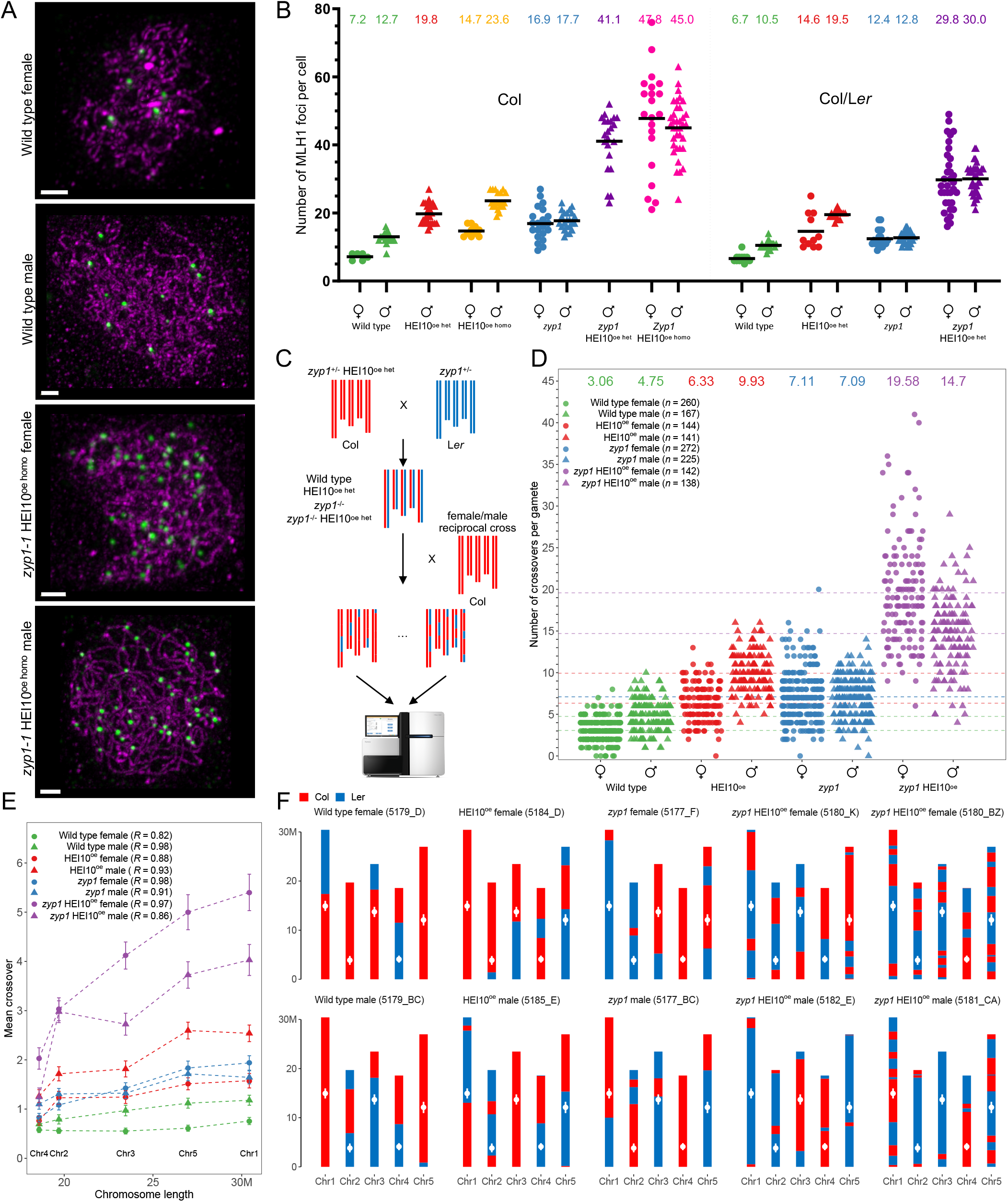
Massive increase in crossovers through combination of *zyp1* mutation and HEI10 overexpression. (A) MLH1 foci in Col wild type and *zyp1* HEI10^oe homo^ meiocytes. Following immunolocalization, REC8 (Purple) and HEI10 (not shown) were imaged with STED while MLH1 (green) was imaged with confocal microscopy. The maximum intensity projection is shown. Scale bar=1µm. (B) Corresponding MLH1 foci quantification, in female and male, inbred Col and hybrid Col/L*er*. The HEI10 transgene originates from the C2 line and is either homozygous (HEI10^oe het^) or heterozygous (HEI10^oe homo^). Each dot is an individual cell, and the mean is indicated by a bar and a number on the top. (C) Experimental design for construction of female and male hybrid populations for sequencing. (D) The number of COs per gamete in female and male populations of wild type, HEI10^oe^, *zyp1*, and *zyp1* HEI10^oe^. Each point is a BC1/gamete, and the means are indicated by horizontal dashed lines and numbers on the top. The population size is shown in parentheses. (E) Correlation analysis between mean number of COs per chromosome per gamete and chromosome size (Mb). Error bars are the 90% confidence intervals of the mean. Pearson’s correlation coefficients are shown in parentheses. (F) Chromosomal genotypes are shown for representative gametes for wild type and mutants, and for extreme cases for *zyp1* HEI10^oe^ populations. Centromere positions are indicated by white points.

**Figure 2.**
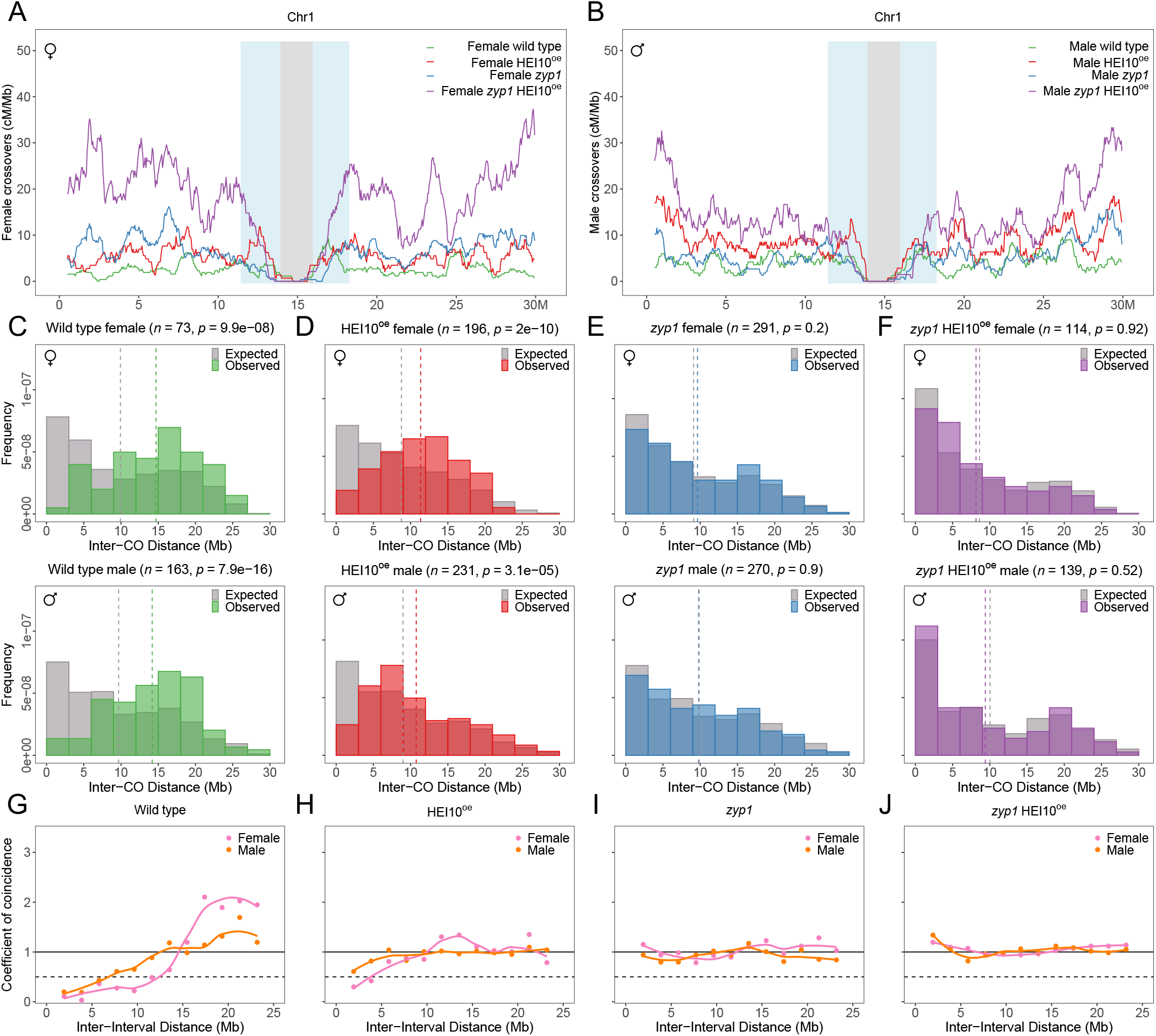
CO distribution and interference analysis in female and male wild type, HEI10^oe^, zyp1, and *zyp1* HEI10^oe^. (A–B) The distribution of COs on chromosome 1 in (A) female and (B) male of wild type, HEI10^oe^, *zyp1*, and *zyp1* HEI10^oe^. The other chromosomes are shown in Figure S3. (C–F) Distribution of distances between two COs for chromosomes with exactly two COs (Figure S5). The grey bar represents the expected distribution of COs without interference, calculated by permutation analysis of COs. The number of analyzed CO pairs and the p-value from the Mann-Whitney test between the expected and observed are indicated. (G-J) CoC curves in female and male meiosis of wild type, HEI10^oe^, *zyp1*, and *zyp1* HEI10^oe^, respectively. Chromosomes were divided into 13 intervals, for calculating the mean coefficient of coincidence of each pair of intervals.

### Overexpression of HEI10 increases COs but maintains heterochiasmy

In wild types, the number of MLH1 foci is higher in males than females in both the inbreds and the hybrids (ratio male/female=1.8 and 1.6, respectively. Figure 1A–B). Whole-genome sequencing of male-and female-derived hybrid progenies showed that CO numbers detected genetically are higher in male meiosis than in female meiosis (Figure 1D, ratio=1.6, p<0.001), confirming heterochiasmy. The number of MLH1 foci at male meiosis is higher in wild-type Col than in Col/L*er*. Analysis of quantitative trait loci (QTL) in a Col/L*er* population previously revealed that the Col *HEI10* allele is associated with higher recombination levels, suggesting that at least a part of this difference in MLH1 counts can be attributed to a difference in HEI10 activity^23^. In wild-type female, the MLH1 foci numbers are not significantly different between Col and Col/L*er* and close to the minimum of one per chromosome (7.2 and 6.8 foci for five chromosomes). In the presence of a transgene ectopically overexpressing HEI10 (HEI10^oe^ C2 line ^23^, homozygous), the number of MLH1 is increased ∼two-fold in both sexes, in both Col and Col/L*er*. Heterozygosity for the HEI10^oe^ transgene also increases MLH1 foci number, but slightly less than homozygosity, confirming the effect of HEI10 dosage on recombination^6,23^ and suggesting that the level of HEI10 in the C2 line is close to saturation. Importantly, increases provoked by HEI10 dosage modulation are similar in males and females, leading to more MLH1 in males than females (p=0.0001) (Figure 1B). This was confirmed with progeny sequencing in hybrids, which revealed a 2.1-fold increase of COs in HEI10^oe^ female and male, compared with wild type (p < 2.2e-16, Figure 1D-F). The ratio of male vs. female COs is maintained at 1.6 in HEI10^oe^ (p < 2.2e-16). In summary, overexpressing HEI10 provokes a doubling of class I COs in both female and male, maintaining heterochiasmy.

### *ZYP1* mutation increases COs and abolishes heterochiasmy

Mutating the transverse element of the SC ZYP1 also increases MLH1 foci number (Figure 1B). In Col *zyp1*, compared to wild type, the numbers increased 1.4-fold in males, consistent with previous findings^10,11^, and 2.3-fold in females. In the Col/L*er* hybrid, the numbers increased by 1.2-fold in male and 1.8-fold in females. In contrast to HEI10^oe^, MLH1 foci number is no longer significantly different in males versus females (p>0.6). This is consistent with genetic crossovers detected by sequencing of hybrid progenies with equal numbers observed in female and male gametes and fold increases of 2.3 in females and 1.5 in males compared to wild type (Figure 1D)^11^. The *zyp1* mutation thus leads to an increase in class I COs, which disproportionately affects female meiosis and abolishes heterochiasmy.

### Combining HEI10 overexpression and *zyp1* massively increases class I COs

HEI10^oe^ and *zyp1* increase CO number, but in different ways; while the former maintains heterochiasmy, the latter does not. We thus combined *zyp1* mutation and HEI10^oe^ and analyzed the effects on MLH1 foci numbers (Figure 1A–B). In Col, the number of foci observed in *zyp1* mutants homozygous for the HEI10^oe^ transgene was significantly higher than ever previously reported, reaching 47.8 and 45.0 in females and males, respectively. The female and male MLH1 counts are not significantly different from each other and represent marked 6.7-fold and 3.5-fold increases compared to their respective wild types. In Col *zyp1* males heterozygous for HEI10^oe^, the MLH1 foci count was slightly but significantly lower (41.1, p=0.015) than the homozygous, showing that there is a dynamic range of HEI10 dosage effects on COs. In the hybrid *zyp1* HEI10^oe^, the observed number of MLH1 in females and males was 29.8 and 30.0, not significantly different from each other (p=0.8) but representing a 4.4-and 2.9-fold increase compared to their wild-type controls. This suggests that class I COs are massively increased in *zyp1* mutants overexpressing HEI10. Indeed, progeny sequencing showed that the number of genetic COs in male *zyp1* HEI10^oe^ was dramatically increased compared to wild type, reaching 14.7 CO per gamete (3.1-fold, Mann-Whitney test, p < 2.2e-16, Figure 1D-F), fitting well with the 30 MLH1 foci counted in male meiocytes (Figure 1B). In females, COs were also vastly increased, but intriguingly, to even higher levels than predicted by the number of MLH1 foci (30/2=15), reaching a striking 19.6 COs per female gamete (6.4-fold/wild type, p < 2.2e-16, Figure 1D–F). Together, this shows that combining *zyp1* mutation and HEI10 overexpression cumulatively and massively increases the numbers of class I COs. It also suggests that class II COs may also be increased in female *zyp1* HEI10^oe^. Such an increase in class I COs is unprecedented, and suggests that the central element of the SC and HEI10 levels are two main regulators limiting class I COs.

Looking along chromosomes, *zyp1* and HEI10^oe^ individually or in combination elicit a massive increase in COs along the arms while the peri-centromeres and the Col/L*er* large inversion^24,25^ remained recalcitrant to recombination (Figure 2, Figure S3). At the fine scale, the majority of COs were located in genic regions in both wild type and mutants (Figure S4). This suggests that despite a large increase in CO number, the local preference for CO placement is conserved, presumably because the distribution of double-strand breaks is maintained. For all eight hybrid populations, the average observed number of COs is positively correlated with the physical size of chromosomes (Pearson’s correlation coefficients >0.8, Figure 1E). We looked for co-variation of CO frequency between chromosomes within the same meiocyte/gamete, as observed in various species^26^. No significant correlation was seen in any of the populations, with maximum correlation coefficients of ∼0.2 observed in female *zyp1* HEI10^oe^ (Figure S2), suggesting that this co-variation does not exist in Arabidopsis or is too small to be detected in our essay.

### CO interference is reduced by HEI10 overexpression and abolished in *zyp1*

To measure the impact of *zyp1* and HEI10^oe^ on CO interference, we first analyzed the distribution of distances between two genetically detected COs for chromosomes with exactly two COs in female and male gametes (Figure 2). In wild-type females and males, the distribution was significantly shifted to large inter-CO distances (p<10^−6^) compared with the expected distribution if the COs were randomly spaced, showing the presence of CO interference (Figure 2C). In HEI10^oe^ females and males, the distribution was also shifted to longer distances, showing the presence of CO interference in both sexes (p<10^−4^, Figure 2D). However, the shift was less marked than in the wild type, suggesting a reduction of interference in HEI10^oe^. In *zyp1* and *zyp1* HEI10^oe^, the observed distributions of inter-CO distances were not different from what would be expected in the case of random spacing (p>0.2, Figure 2E– F), suggesting an abolition of CO interference in both females and males. Furthermore, we performed a coefficient of coincidence (CoC curve) analysis for an alternative, likely more accurate, measurement of CO interference (Figure 2G-J)^9^. In wild type, the two CoC curves are below 1 at distances <∼15 Mb in both females and males, confirming the presence of substantial CO interference (Figure 2G). The female curve stays close to 0 for longer distances, showing that CO interference propagates to longer Mb distances in females, consistent with previous analyses^11,16,18^. In HEI10^oe^, the curves also deviate from 1 at short distances (<∼7Mb), showing the presence of interference, although at a reduced level compared to wild type (Figure 2H). As in wild type, interference in HEI10^oe^ is stronger in female than in male meiosis. In contrast, the CoC curves are flat at values close to 1 for both females and males in *zyp1* (Figure 2I), confirming that CO interference is abolished in the absence of ZYP1^10,11^. In *zyp1* HEI10^oe^, the curves are also flat at ∼1, showing that the numerous class I COs produced in this context do not interfere with each other (Figure 2J). Thus, HEI10^oe^ reduces, while *zyp1* abolishes CO interference.

### High CO rates in *zyp1* and HEI10^oe^ are not associated with meiotic defects

The limited level of COs per chromosome observed in most eukaryotes could suggest that a high level of COs has a detrimental effect. We explored if a massive elevation of class I COs is associated with meiotic chromosome segregation and fertility defects. The number of seeds per fruit is reduced in *zyp1-1* compared to wild type (−8%, t-test p<0.001), consistent with previous results and the reported loss of the obligate CO in *zyp1* mutants^10,11^ (Figure 3I). Analyses based on sequence coverage detected a few aneuploids among *zyp1* gametes (2/497, Figure 3J and S6–7) that were not detected in hybrid wild types (0/427 in this study, and 0/760 in an independent wild-type dataset^27^). The HEI10^oe^ C2 line also showed a slight reduction of fertility (−12%, p=0.005, Figure 3I) and low frequency of aneuploid gametes (2/285). In *zyp1* HEI10^oe^, seed number was reduced (−7%, p=0.025), and a small number of aneuploids were detected in hybrids (7/272), suggesting a slight meiotic defect also in this background. All the 11 identified trisomy cases concerned chromosome 4, the shortest Arabidopsis chromosome. The centromeric region of chromosome 4 of the aneuploid gamete is systematically heterozygous Col/L*er*, which is diagnostic for missegregation at meiosis I (failure to separate homologous chromosomes). For the vast majority (9/11), no CO was detected on the aneuploid chromosome, which is compatible with an absence of COs in the bivalent. This suggests that these nine events resulted from the loss of the obligate crossover and consequent random missegregation of homologs. Two aneuploids, both from *zyp1* HEI10^oe^, had two COs on the trisomic chromosome. In both cases, the two COs are relatively close to each other (∼2 and 4 Mb), which may lead to an unstable connection between the homologs as spindle tension would be counteracted by only a short stretch of cohesion. The aneuploidies appear thus to be associated with the absence of COs or specific configurations of a pair of COs. Meiotic chromosome spreads in *zyp1* HEI10^oe^ showed that most metaphase I cells had a wild-type configuration with five bivalents aligned on the metaphase plate (44/45 in Col; 23/30 in Col/L*er*; Figure 3C). However, one univalent was observed in a minority of cells (1/45 and 7/30, Figure 3C). Consistently, at metaphase II almost all cells had five chromosomes aligned on the two plates (25/25 and 6/7; Figure 3D), and one had a 6:4 configuration indicating unbalanced segregation at meiosis I (Figure E, F), likely due to the absence of the obligate CO. Altogether, this shows that a slight meiotic chromosome segregation defect is present in HEI10^oe^ *zyp1*. However, the rare missegregations appear to be due to an incomplete CO assurance and are not associated with the extreme CO numbers observed in the mutants (up to 15 COs in a single chromatid, Table S4). This suggests that high CO number does not impair chromosome segregation and raises the question of the evolutionary forces that limit CO to typically less than three per chromosome per meiosis in most eukaryotes^1,28^. While failure to insure at least one CO per chromosome pair is associated with meiotic failure in most eukaryotes, the reasons that prevent high CO number are unclear. The absence of an immediate cost of massively elevated CO numbers in HEI10^oe^ *zyp1* suggests that low CO numbers are not selected for by evolution because of mechanical constraints during meiosis. Rather, this observation suggests that the medium-to-long term genetic effects of COs are subject to indirect selection^1^. This supports the suggestion that a relatively low recombination rate, not much higher than one per chromosome, is optimal for adaptation.

**Figure 3.**
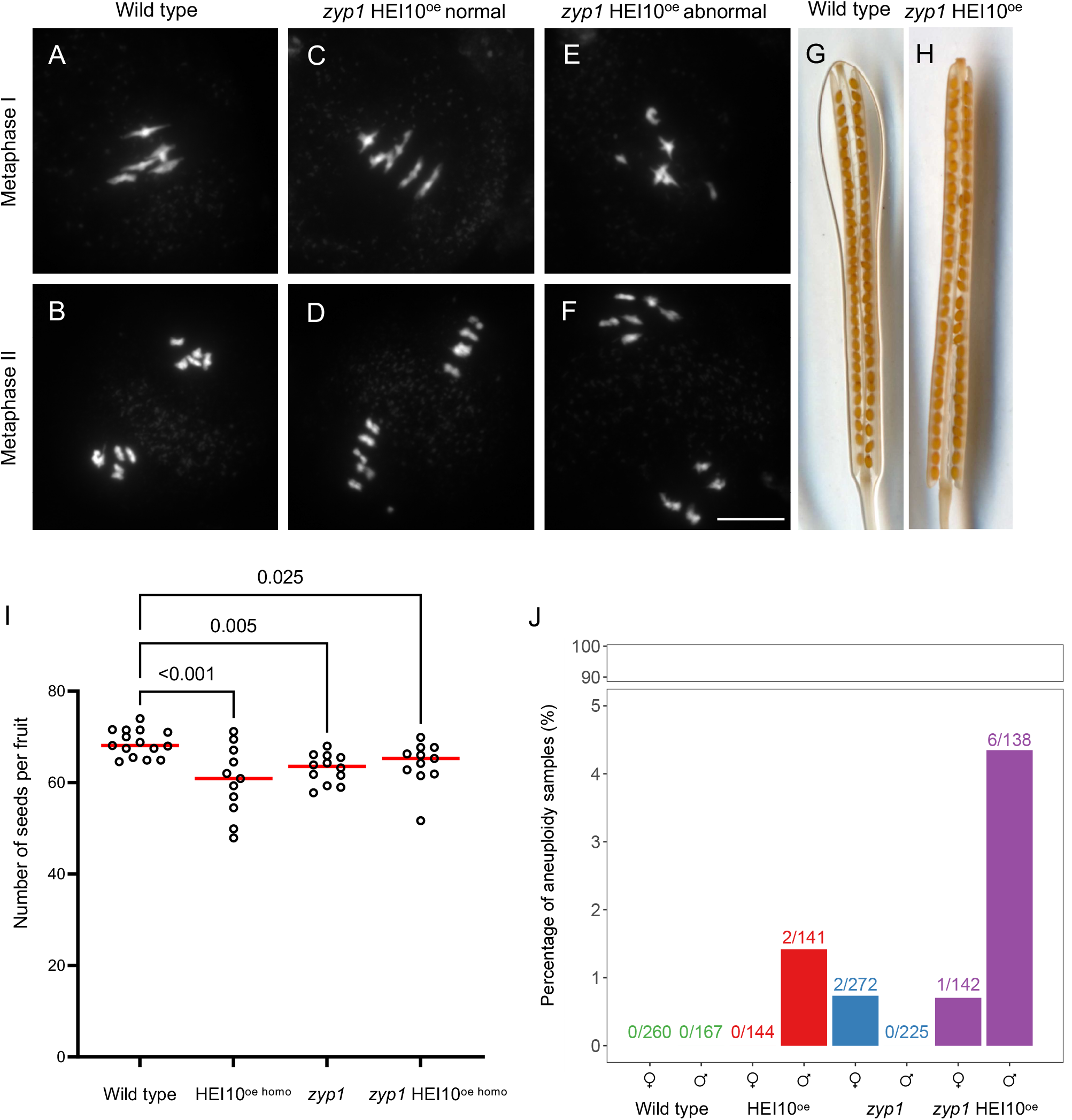
Analysis of meiotic and fertility defects. (A–F) DAPI-stained meiotic chromosome spreads from Col/L*er* male meiocytes in wild type (A, B) and *zyp1* HEI10^oe^ (C-F). (A, C, E) Metaphase I. (B, D, F) Metaphase II. (C, D) Normal chromosome configurations in *zyp1* HEI10^oe^. (E, F) Rare abnormal chromosome configurations in *zyp1* HEI10^oe^. Scale bar=10µm. (G–H) Representative cleared fruits of wild-type Col and *zyp1* HEI10^oe^ mutants. (I) Corresponding quantification of fertility. Each dot represents the fertility of an individual plant, measured as the number of seeds per fruits averaged on ten fruits. The red bar shows the mean. All plants were siblings grown together in a growth chamber. *P* values are one-way ANOVA followed by Fisher’s LSD test. (J) The percentage of aneuploid samples detected in each population (Figure S6-7). The proportion of aneuploid samples in each population is shown on top of the bars.

### Female and male SC lengths differ and are affected by neither HEI10^oe^ nor *zyp1*

SC length has been shown to correlate with the frequency of class I COs^4,16,17^. We wondered if the class I CO increase provoked by *zyp1* and HEI10^oe^ is associated with variation in SC length. We traced chromosome axis (REC8) in female and male meiocytes with preserved 3D organization and measured the length of each chromosome (Figure 4, Table S5). In wild type, we found that the SC is 1.6-fold longer in males than females, consistent with previous reports^16^ (Figure 4M). The longer total SC length in wild-type males is proportional to the higher MLH1 foci and CO numbers compared to females (Figure 1B, 1D, 4Q), suggesting that SC length determines CO number and thus drives heterochiasmy. Strikingly SC/axis absolute and relative length is conserved in both sexes in HEI10^oe^, *zyp1*, and *zyp1* HEI10^oe^ mutants, thus maintaining the male-female dimorphism (Figure 4M). In HEI10^oe^, the MLH1 foci and CO numbers are increased proportionally in males and females, maintaining heterochiasmy (Figure 1B, 4O–Q). This suggests that the effect of HEI10 dosage on COs is constrained by the length of the SC. In clear contrast to HEI10^oe^, the link between axis length and CO number is disrupted in *zyp1*, with MLH1 foci and COs equal in males and females despite a large difference in axis length (4O–Q). The observation that the length of pairs of axes in *zyp1* matches the length of the assembled SC in the wild type suggests that the length of the two axes directly determines SC length. In the double mutant *zyp1* HEI10^oe^, MLH1 foci are massively increased and reach equal numbers in males and females despite different axis lengths that are unmodified compared to wild type (Figure 4P). This suggests that HEI10 dosage has a comparable effect in males and females in the absence of the SC.

**Figure 4.**
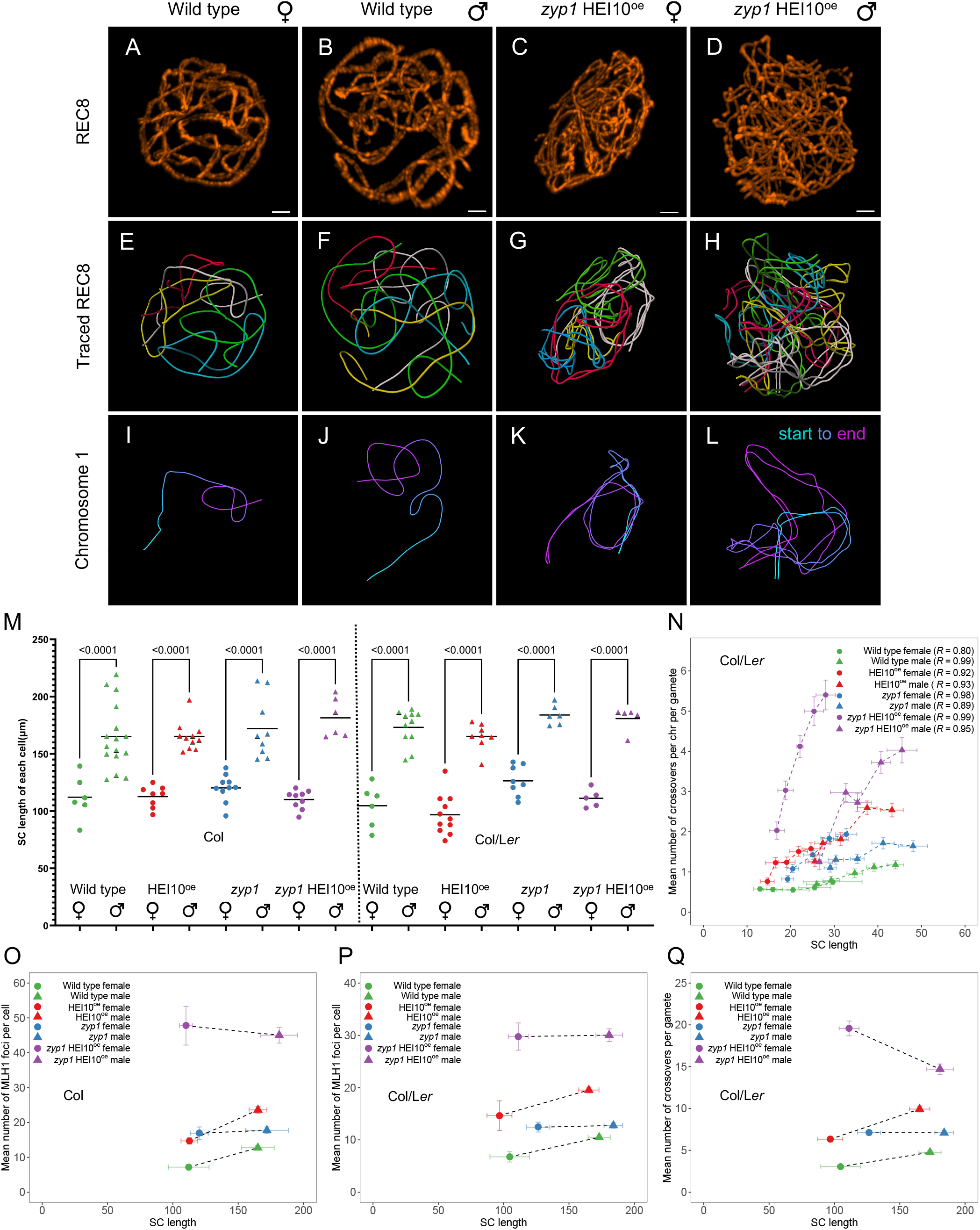
Analysis of SC/axis lengths in female and male meiocytes. (A–D) REC8 immunolocalization in female and male meiocytes of wild type and *zyp1* HEI10^oe homo^ (Col). Imaging was done with 3D-STED and the projection is shown. Scale bar=1µm. (E– H) REC8 signal was traced in 3D. Each bivalent pair is color-coded. (I–L) Individual trace of the longest chromosome (presumably chromosome 1), with start-to-end color code. (M) Measurement of the total SC length. Each dot is the SC length of an individual cell. The bars indicate the mean. One-way ANOVA followed by Sidak correction showed that SCs were systematically longer in males than in females (p < 0.0001). The same test did not detect any differences between any of the pairs of males of different genotypes (p>0.4). For females, none of the pairwise comparisons were significantly different except in Col/L*er* HEI10^oe^ that was lower than Col/L*er zyp1* (p=0.006) and Col *zyp1* (p=0.008). Note that variations in slide preparation and exact meiotic stage may affect this result. (N) Correlation analysis between the mean number of COs per chromosome per gamete and SC length (µm) in Col/L*er* background. SCs were attributed to specific chromosomes based on their length (e.g., the longest was presumably chromosome 1). Pearson’s correlation coefficients are shown in parentheses. (O– P) The relationship between the mean number of MLH1 foci per cell and total SC length per cell in (O) Col background and (P) Col/L*er* background. (Q) The relationship between the mean number of COs per gamete and SC length in Col/L*er* background. The 90% confidence intervals are indicated as error bars.

Altogether, this suggests that two major factors conjointly regulate CO number: (i) Our results show that the central element of the SC ZYP1 imposes interference and limits COs. The length of the axis/SC is correlated with the number of COs in various contexts^4^, and notably when comparing sexes. Crucially, this correlation is lost in the absence of ZYP1, where the difference in axis length is no longer associated with a difference in CO number, suggesting that COs are regulated by the length of the tripartite SC and thus indirectly by the axis. The upstream mechanisms that determine the differences in SC lengths in males and females in many organisms remain to be determined. (ii) HEI10 dosage positively regulated CO formation. The effect of HEI10 dosage appears to be constrained by the length of the SC. HEI10 initially loads as multiple foci along the SC before consolidating into a small number of large foci at CO sites^19^. This supports a model in which HEI10 loading on the SC depends conjointly on HEI10 expression levels and SC length and that this loading eventually determines CO number.

### The HEI10 coarsening model

The results we present here and previous observations can be interpreted in the context of an emerging model for crossover patterning via droplet coarsening through the diffusion of HEI10 along the SC^5,6^. In this model (Figure 5), HEI10 initially forms multiple droplets along the SC, and HEI10 molecules diffuse along the SC from droplets to droplets. If larger droplets tend to retain more HEI10 molecules than smaller droplets, a coarsening process is initiated, and large droplets grow at the expense of nearby smaller droplets, leading to the formation of well-spaced large droplets. These large droplets are proposed to create a specific context that promotes class I CO formation (e.g., by attracting the MLH1/MLH3 complex) and protects recombination intermediates from anti-CO factors (i.e., FANCM and RECQ4^29,30^). It is unclear if initial droplets colocalize with recombination intermediates or if recombination intermediates favor the coarsening process locally, but both hypotheses envisage final droplets to embed such an intermediate. This model predicts the obligate crossover, a limited number of COs, and interference^6^. If the coarsening process can proceed without restrictions, it would ultimately lead to the formation of a single droplet/CO per bivalent, as observed in *C. elegans*^*5*^. However, in most species, including Arabidopsis, 2–3 interfering class I COs are typically observed per bivalent. At least three hypotheses can account for this observation: one proposes an upper limit in the size of a droplet, above which it stops growing, allowing other droplets to be maintained. The second supposes that the coarsening is stopped when a checkpoint is satisfied (e.g., when a least one large droplet/one CO is formed per chromosome). The third suggests that the process is stopped before completion after a certain period, which we consider here for simplicity. In all cases, the total amount of HEI10 loaded onto the SC determines the number of CO-promoting droplets, although in the third case the length of the SC also plays a minor role independently of the total amount of HEI10. The model proposes that two factors jointly determine the initial HEI10 loading: (i) HEI10 concentration in the nucleoplasm, which determines the amount of HEI10 in initial droplets and on the SC per µm of SC (ii) the length of the SC, which, for a given expression level of HEI10 would determine linearly the total HEI10 loading. Our numerical implementation of this model (see Methods) explains the measured CO counts quantitatively (Figure 6A,B). In particular, it explains the observed correlation between the length of the SC and the number of COs between chromosome pairs within single cells as well as between different cells, as notably observed here in Arabidopsis, where female meiosis has a shorter SC and fewer COs than male meiosis. Note that this shorter SC in females also implies stronger CO interference (Figure 6C,F, G; compare with Figure 2). This model also accounts for the fact that CO number depends on HEI10 expression level, as this level determines the amount of HEI10 loaded per µm of SC. Remarkably, we observed that overexpressing HEI10 increases CO numbers in males and females without eliminating heterochiasmy, as predicted by the difference in SC length. In addition, CO interference is also reduced, but not abolished by over-expressing HEI10, as expected, as the coarsening process still occurs (Figure 6D,F,H, I). We propose that in the absence of SC, in the *zyp1* mutant, HEI10 diffusion is no longer constrained to the SC but occurs freely in the nucleoplasm. In this case, droplets still form on chromosomes (Figure 1A-B), but they now exchange HEI10 directly with the nucleoplasm. If this exchange is slow compared to the duration of pachytene, all initial droplets grow continuously by taking up HEI10. In contrast, when HEI10 is exchanged more quickly, competition between droplets, and thus coarsening, will set in, which was also recently proposed^31^. In both cases, large HEI10 foci form, colocalize with MLH1, and promote class I COs. However, the obligate CO and CO interference are lost as the diffusion is no longer constrained per chromosome (Figure 6E,J). In a sense, in the absence of the SC, the coarsening and CO designation process can be said to be “blind” to chromosomes. The absence of the SC must be associated with slower coarsening since otherwise the exchange of HEI10 via the nucleoplasm would be significant in wild type, too. If the number of initial droplets in the *zyp1* mutant is roughly comparable to wild type, slower coarsening implies a bigger number of large droplets at the end of pachytene, consistent with the increase observed experimentally (Figure 1). Together with interference, heterochiasmy is abolished when the number of COs per chromosome is solely determined by HEI10 expression level in the nucleoplasm and no longer by HEI10 loading onto the SC. Taken together, the experimental data and the coarsening model show that two factors limit class I COs: ZYP1-mediated CO-interference and HEI10 levels.

**Figure 5.**
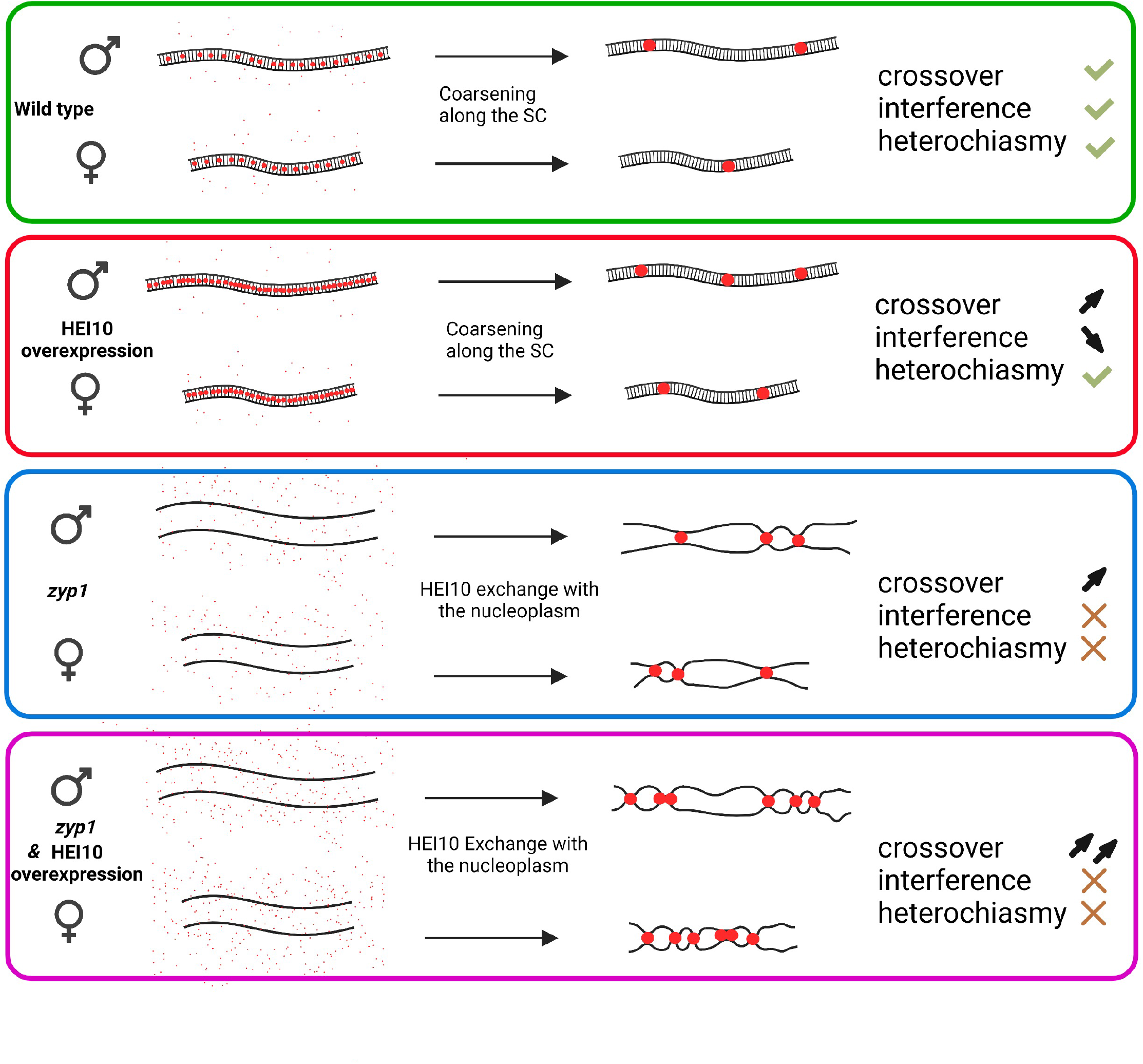
Model of crossover patterning *via* HEI10 coarsening. HEI10 (red) is captured by the central element of the SC and coarsens into large pro-CO droplets. The number of large pro-CO droplets is determined by SC length (heterochiasmy), and HEI10 expression levels. HEI10 overexpression increases CO number, and weakens interference but maintains heterochiasmy. In absence of an SC (*zyp1*), HEI10 is exchanged directly between the droplets and the nucleoplasm abolishing both interference and heterochiasmy, and the number of droplets depends on HEI10 expression level. Created with BioRender.com

**Figure 6.**
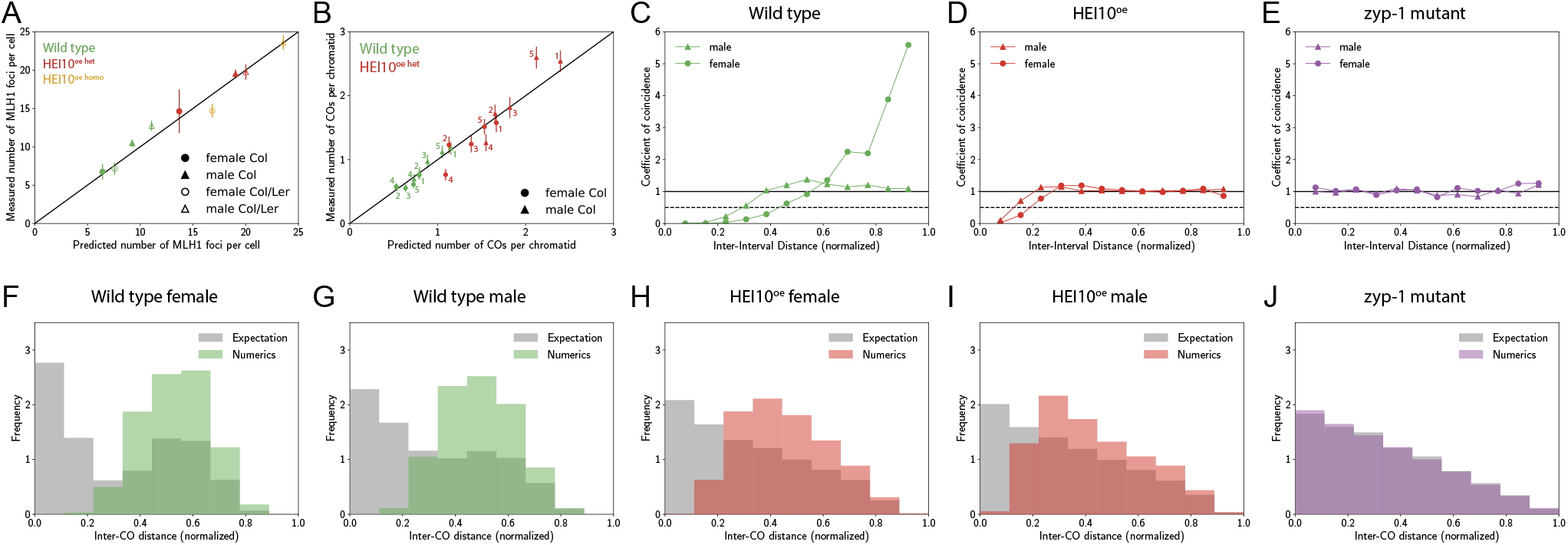
A coarsening model for crossover designation explains the measured data. (A) Number of MLH1 foci predicted by the model compared to the experimental measurements shown in Figure 1B. Error bars denote 90% confidence. (B) Number of COs per chromatid predicted by our model compared to the experimental measurements shown in Figure 1D. The respective chromatids are labeled and error bars denote 90% confidence. (C–E) Predicted coefficient of coincidence curves; compare to Figure 2G–J. (F–J) Predicted distributions of distances between two COs for chromosomes with exactly two COs; compare to Figure 2C–F. (A–J) Numerical details are given in the Methods. Mean and confidence intervals were determined from *n* = 1000 (except *n* = 500 in panel A) independent repetitions.

A similar model was proposed and further supporting experimental data were recently obtained in *C. elegans*^5^. Several additional pieces of evidences suggest that the dual control of COs by SC and HEI10 is conserved: In multiple species, HEI10 homologs also initially form multiple foci before eventually consolidating into a limited number of large foci that co-localize with COs^19-22,32^; COs covary with SC length in many species^4^; Variants that affect recombination rates in natural populations of diverse species involve genes that encode HEI10 homologues^33^. This suggests that the coarsening of HEI10 along the SC may be a conserved process for CO patterning in eukaryotes.

## Materials and methods

### Plant Materials and Growth Conditions

*Arabidopsis thaliana* plants were cultivated in Polyklima growth chambers (16-h day, 21.5 °C, 280 µM; 8-h night, 18 °C: 60% humidity). Wild-type Col-0 and L*er*-1 are 186AV1B4 and 213AV1B1 from the Versailles stock center (http://publiclines.versailles.inra.fr/). The *zyp1-1* (8.7.2V1T3) and *zyp1-6* (1.12V5T2) mutants were previously described^11^. The HEI10 over-expression line is Col HEI10 line C2^23^, kindly provided by Ian Henderson. Genotyping of the mutants was carried out by PCR amplification (Dataset S1).

To generate the double homozygous mutant *zyp1-1*^*-/-*^ HEI10^oe^ in Col, *zyp1-1*^*+/-*^ plants were crossed with HEI10^oe^ homozygous mutant plants (C2). The obtained double heterozygous *zyp1-1*^*+/-*^ HEI10^oe^ were selfed to produce *zyp1-1*^*-/-*^ mutants, HEI10^oe^ homozygous, and *zyp1-1*^*-/-*^ HEI10^oe^ double homozygous mutants. These sister plants were used to perform MLH1 foci counting, SC measurements, chromosome spreads, and seed countings. To generate *zyp1-1/zyp1-6* HEI10^oe het^ in Col/L*er*, double heterozygous *zyp1-1*^*+/-*^ HEI10^oe^ (Col) were crossed with *zyp1-6*^*+/-*^ (L*er*) to generate *zyp1-1/zyp1-6* HEI10^oe het^, HEI10^oe het^, *zyp1-1/zyp1-6* and wild-type controls in Col/L*er*. These sister plants were used for MLH1 foci counting and SC length measurements and were reciprocally backcrossed with wild-type Col to generate the sequencing populations. Backcross populations were grown in the greenhouse for three weeks (16-h day/8-h night) and four days in the dark. For DNA extraction and library preparation, 100–150mg leaf samples were collected from the four backcross populations ^34^.

### Cytology

Chromosome spreads and immunolocalization of male meiocytes on 3D slides were conducted as previously described^35,11^. For female 3D slides, 0.8–1.2mm pistils were collected and their stigmata cut off. Pistils were then fixed and digested following the same procedure as for male meiocytes. The pistils were then opened longitudinally and the ovules released on a slide. The subsequent slide treatment and immunolocalization were the same as for male meiocytes, and were described previously^11^.

Four primary antibodies were used: anti-REC8 raised in rat^36^ (laboratory code PAK036, dilution 1:250), anti-MLH1 in rabbit^37^ (PAK017, 1:200), anti-HEI10 in chicken^19^ (PAK046, 1:5,000) and anti-ZYP1N in guinea pig^11^ (PAK053, 1:500). Secondary antibodies were STAR RED, STAR ORANGE and STAR GREEN, or Alexa488. Super-resolution images were acquired with the Abberior instrument facility line (https://abberior-instruments.com/) 561-and 640-nm excitation lasers (for STAR Orange and STAR Red, respectively) and a 775-nm STED depletion laser. Confocal images were taken with the same instrument with a 485-nm excitation laser (for STAR GREEN/Alexa488).

### Image processing and analysis

Deconvolution of the images was performed by Huygens Essential (version 21.10, Scientific Volume Imaging, https://svi.nl/) using the classic maximum likelihood estimation algorithm with lateral drift stabilization; signal-to-noise ratio: 7 for STED images and 20 for confocal images, 40 iterations, and quality threshold of 0.5. Maximum intensity projections and contrast adjustments were also done with Huygens Essential. Deconvoluted pictures were imported into Imaris 9.8 (https://imaris.oxinst.com/, Oxford Instruments, UK) for subsequent analysis. MLH1 foci were counted using the spots module in diplotene and diakinesis cells. The vast majority of MLH foci colocalize with a HEI10 focus. Only double MLH1/HEI10 foci present on chromosomes were taken into account. For REC8 signal tracing, fully synapsed cells were used to trace the chromosomes. In wild type and HEI10^oe^, the five synapsed bivalents were traced. In *zyp1* and *zyp1* HEI10^oe^, five pairs of parallel chromosomes were traced. The surface module was used to create a clean masked REC8 channel for filament tracing. The filament module was used to trace the SC length, AutoDepth function was used to do semi-automatic tracing and get the simulated chromosome. The SC length of each chromosome was measured using the statistics function of the Filament module.

### CO identification and analysis

In this study, the female and male population of wild type (48 and 47 plants), HEI10^oe^ (144 and 141 plants), *zyp1* (48 and 47 plants) and *zyp1* HEI10^oe^ (142 and 138 plants) were sequenced by Illumina HiSeq3000 (2×150bp) conducted by the Max Planck-Genome-center (https://mpgc.mpipz.mpg.de/home/). The raw sequencing data of the female and male population of wild type (212 and 120 plants respectively) and *zyp1* (224 and 178 plants) from a previous study (ArrayExpress number E-MTAB-9593) ^11^ were also included in this study. In total, we analyzed 260 and 167 wild type female and male, 144 and 141 HEI10^oe^ female and male, 272 and 225 *zyp1* female and male, 142 and 138 *zyp1* HEI10^oe^ female and male plants, separately. The raw sequencing data were quality-controlled using FastQC v0.11.9 (http://www.bioinformatics.babraham.ac.uk/projects/fastqc/). The sequencing reads were aligned to the *Arabidopsis thaliana* Col-0 TAIR10 reference genome, which was downloaded from TAIR ^38,39^, using BWA v0.7.15-r1140^40^, with default parameters. A set of Sambamba v0.6.8^41^ commands was used for sorting and removing duplicated mapped reads. The creation of the high-confidence SNP marker list between Col and L*er*, meiotic CO detection (a sliding window-based method, with a window size of 50 kb and a step size of 25 kb), check and filtering of low covered and potential contaminated samples were performed according to previous protocols^11,27,42-44^. Samples of each population were randomly selected for checking predicted COs manually by inGAP-family^43^. The Coefficient of Coincidence (CoC) was calculated for CO interference analysis using MADpattern^45,46^, with a number of 13 intervals. Chromosome 4 was excluded from interference analyses. To profile the CO distribution along chromosomes, CO position was defined randomly in the range of CO interval and a sliding window-based strategy was used, with 1 Mb window size and 50 kb step size. Then, the local distribution of recombination (CO resolution <= 1000 bp) was explored by ChIPseeker v1.22.1^47^, with the promoter region defined as 2000 bp upstream of the transcription start site.

### Aneuploidy screening by whole genome sequencing

The sequencing depth of each non-overlapping 100 kb window across the genome was evaluated by Mosdepth v0.2.7^48^ with parameters of “-n --fast-mode --by 10000”. For each sample, pairwise testing of sequencing depths along chromosomes was performed using the Mann-Whitney test, and significant *p* values were adjusted using the fdr method. A pair of tested chromosomes with fold change > 1.2 and *p* value < 1e-20 was considered as aneuploid.

### Mathematical model of CO patterning

Our mathematical model describes the concentration *c*(*x, t*) of HEI10 along the SC of length *L* together with the amounts *M*_*i*_(*t*) of HEI10 in N droplets that are placed at positions *x*_*i*_ along the SC for *i* = 1, …, *N*. Here, *x* denotes the length along the SC and t denotes time. Similar to the model presented in ref.^6^, we account for the diffusion of HEI10 along the SC and the exchange of HEI10 between SC and droplets. Droplet *i* grows if the local HEI10 concentration on the SC, *c*(*x*_1_), is larger than the equilibrium concentration 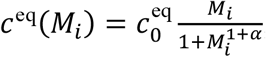

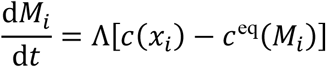

where Λ quantifies the rate of HEI10 exchange. HEI10 diffuses with diffusivity D along the SC and is exchanged with droplets,

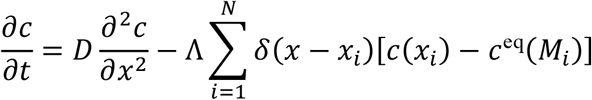

We impose no-flux boundary conditions at *x*= 0 and *x*=*L*, so the total amount of HEI10 is conserved. We implemented this model using finite differences by discretizing the SC using 50 grid points and solved the resulting equations using an explicit Euler scheme.

We initialize the system with a uniform concentration on the SC, *c*(*x, t*= 0) = *c*_init_. The *N* droplets are positioned uniformly along the SC and their sizes *M*_*i*_ are chosen independently from a normal distribution with mean M_init_ and standard deviation *σ*, which has been truncated to [M_init_ - 3*σ*, M_init_ + 3*σ*]. The diffusivity *D* = 1.1 µm^2^/s, the exchange rate Λ= 2.1 µm/s, the exponent *α* = 0.25, and the base equilibrium concentration 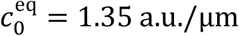, are inspired by ref.^6^. We use SC lengths *L* measured in wild type (Figure 4N) and estimate an initial droplet density of four droplets per µm, based on cytology^49^. For simulations, we choose M_init_ = *y* 3.4 a.u., *σ*_init_ = *y* 1.1 a.u., and *c*_init_ = *y* 1.4 a.u./µm, where *y* is a factor to account for higher HEI10 expression levels. We chose *y*= 2 for wild type Col,*y* = 6 for HEI10^oe het^ Col, *y*= 8 for HEI10^oe homo^ Col, *y* = 1.5 for wild type Col/L*er*, and *y* = 5.5 for HEI10^oe het^ Col/L*er*, which accounts for the reduced activity in L*er*^50^ and HEI10 overexpression. We simulate droplet coarsening on each individual SC for male and female meiosis for 10h, comparable to the duration of pachytene^51,52^. Only droplets above a threshold size of *M*_thresh_ = 3 a.u. are assumed to attract MLH1 and form class I COs. The associated COs per chromatid were determine by choosing COs from the bivalent independently with 50% probability.

In the case of the *zyp1* mutant, our model implies that all CO positions are independent. To obtain a theoretical distribution of COs, we thus first determine the number of COs per chromatid by sampling a Poisson distribution with a mean given by the experimental data (Figure 4N) and then distribute these COs uniformly along the chromatid length.

## Data Availability

MLH1 counts and SC length measurements are shown in table S1 and S6, respectively. The raw sequencing data is deposited in ArrayExpress of EMBL-EBI with accession numbers E-MTAB-11696. The list of identified COs in the female and male populations of wild type, HEI10^oe^, *zyp1*, and *zyp1* HEI10^oe^ can be accessed in Supplemental Table S2-S5.

## Acknowledgements

This work was supported by core funding from the Max Planck Society to R.M., M.E., as well as D.Z. and an Alexander von Humboldt Fellowship to Q.L. and J.J. The IJPB benefits from the support of Saclay Plant Sciences-SPS (ANR-17-EUR-0007). We thank Abby Dernburg for enlightening discussions. We thank Ian Henderson for kindly providing the C2 line. We thank Neysan Donnelly for proofreading the manuscript.

## Author contributions

S.D. produced and analyzed the MLH1-HEI10 cytological data, developed the protocol for female immunolocalization, produced all the genetic material, and analyzed fertility. Q.L. analyzed the sequencing data and performed recombination, interference, and aneuploidy analyses. J.J. generated SC images, and analyzed chromosome missegregation and SC length data. M.E. and D.Z. developed the mathematical model of CO patterning. M.G. developed the method for chromosome 3D analyses. R.M. lead the project and wrote the manuscript with input from all co-authors.

Supplementary Information is available for this paper

Correspondence and requests for materials should be addressed to Raphael Mercier

## Supplemental Figure legends

**Figure S1. Comparison of CO numbers between previous studies and this work**.

The number of COs per gamete in female and male populations of wild type and *zyp1*, respectively. Different genotypes and studies are indicated through different colors and shapes, respectively. The mean CO number of the population and significant *p* value (Mann-Whitney test) are indicated in the top parentheses, with the same color codes.

**Figure S2. Correlation analysis of CO numbers between chromosomes**.

Pearson’s correlation analysis of CO numbers between chromosomes in the same gamete was performed in female and male populations of wild type, HEI10^oe^, *zyp1*, and *zyp1* HEI10^oe^ individually. The sum of COs detected on chromosomes 1 and 2 was plotted against the sum of COs on chromosomes 3 and 5 in the same gamete. A jitter function was applied to avoid points overlapping. The correlation coefficients are shown in parentheses. The very low correlation is in contrast with observations in several other species^26^ and may be due to lower cell-to-cell variation in Arabidopsis.

**Figure S3. The distribution of CO frequency along chromosomes**.

Comparison of CO distributions (sliding window-based, with window size of 1 Mb and step size of 50 kb) between female and male of wild type (A), HEI10^oe^ (B), *zyp1* (C) and *zyp1* HEI10^oe^ (D). Comparison of CO distribution among populations in female (E) and male (F) meiosis. The pericentromeric and centromeric regions are indicated by grey and blue shading, respectively. Consistent with previous observations^11,53^, there is a ∼2.2 Mb region on the long arm of chromosome 4 where recombination is suppressed, which suggests a structural arrangement between the Col and L*er* strains.

**Figure S4. The fine-scale distribution of COs**.

(A) The distribution of the length of CO intervals across populations. The length of 1 kb and 2 kb are indicated by grey dashed lines. The number of analyzed COs is shown in parentheses. The median of CO intervals is 819 bp (B) The distribution of proportion of COs with interval lengths less than 1 kb (high-resolution), more than 2 kb, and the rest separately. (C) The distribution of proportion of high-resolution COs overlapping with genomic features. The promoter region is defined as the 2 kb upstream of the transcription start site. The proportion of the different genomic features is shown as the bar on the top, which is defined by following the priority of promoter, 5’ UTR, 3’ UTR, exon, intron, and intergenic regions. (D) The distribution of distance of high-resolution COs from nearest TSS.

**Figure S5. The distribution of positions of double-COs**.

Relative position of COs for chromosomes with exactly two COs (as in figure 2C-F). The position of the first and second CO of the pair, in female and male meiosis of wild type, HEI10^oe^, *zyp1*, and *zyp1* HEI10^oe^, respectively.

**Figure S6. The analysis of sequencing depths along chromosomes for aneuploidy screening**.

The sequencing depth was calculated for each 100 kb non-overlapped interval along chromosomes. The Mann-Whitney test was used for checking the differences between pairs of chromosomes, the significant p value was then adjusted by the fdr method. The identity of the detected aneuploidy is shown with the same color codes as for the corresponding populations.

**Figure S7. Sequencing depth along aneuploid chromosomes**.

The sequencing depth was calculated for each 100 kb non-overlapped interval along chromosomes. The pericentromeric and centromeric regions are indicated by grey and blue shading, respectively. The horizontal dashed line indicates the mean sequencing depth of the sample. Aneuploidy is visible by higher coverage of one chromosome compared to the others. The label of the detected aneuploidy and corresponding populations are presented individually.

**Table S1. Raw data of MLH1 counts**.

**Table S2. CO positions in wild-type population.**

**Table S3. CO positions in HEI10oe population.**

**Table S4. CO positions in *zyp1* population**.

**Table S5. CO positions in *zyp1* HEI10oe population.**

**Table S6. Raw data of SC Lengths**.

**Table S7. Summary data for mean MLH1 foci number, CO number, and SC length.**

**Table S8. Genotyping primers**

