## supplemental figures for "Dual control of meiotic crossover patterning"

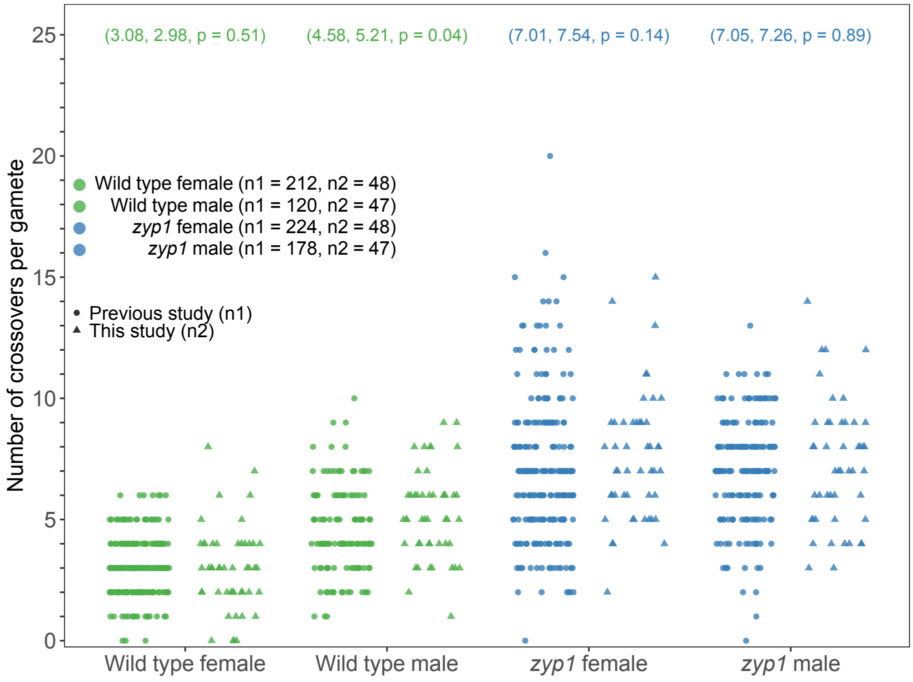


**Figure S1. Comparison of CO numbers between previous and this study..**
The number of COs per gamete in female and male populations of wild type and *zyp1*, respectively. The color-coded and shape-coded points indicate different population and study separately. The mean CO number of the population and significant p value (Mann-Whitney test) are indicated in the top parentheses, with same color codes.

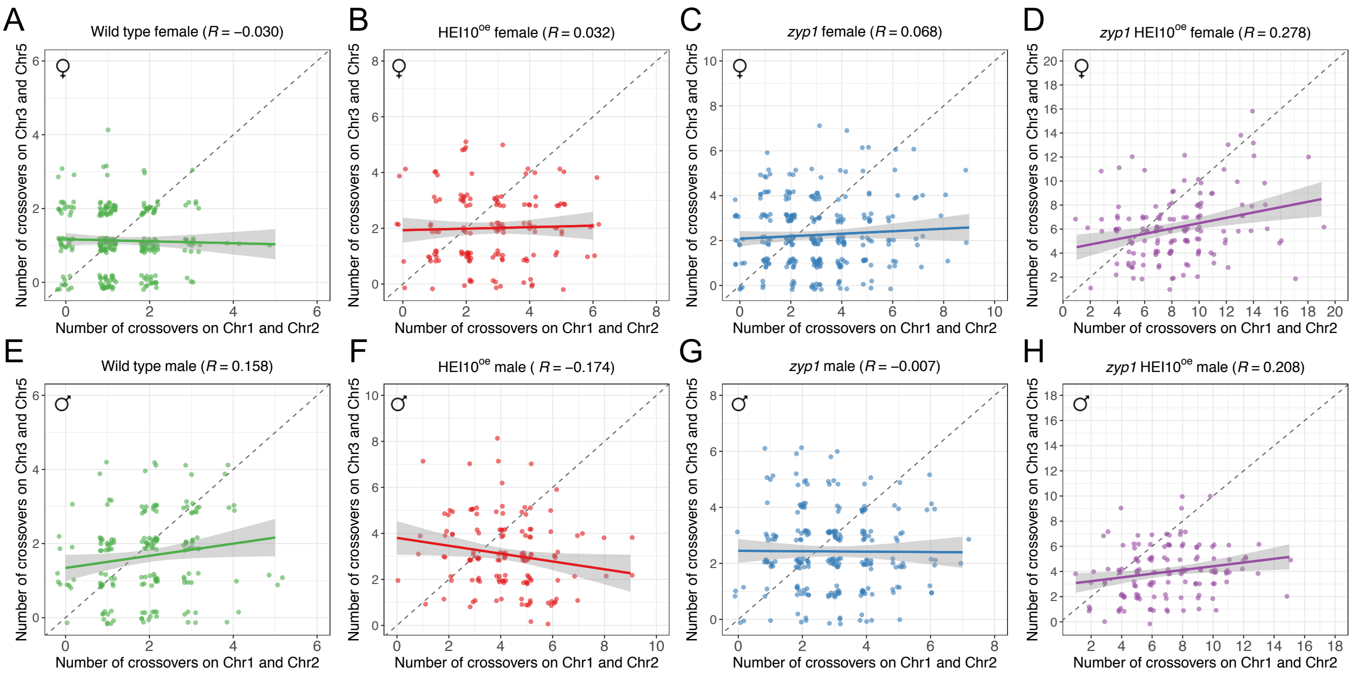


**Figure S2. Correlation analysis of CO numbers between chromosomes.**

Pearson's correlation analysis of CO numbers between chromosomes in the same gamete was performed in female and male populations of wild type, HEI10$^{oe}$, *zyp1* and *zyp1* HEI10$^{oe}$ individually. The sum of COs detected on chromosomes 1 and 2 was plotted against the sum of COs on chromosomes 3 and 5 in the same gamete. A jitter function was applied to avoid points overlapping. The correlation coefficients are shown in parentheses. The very low correlation is in contrast with observations in several other species[26] and may be due to lower cell-to-cell variation in Arabidopsis.

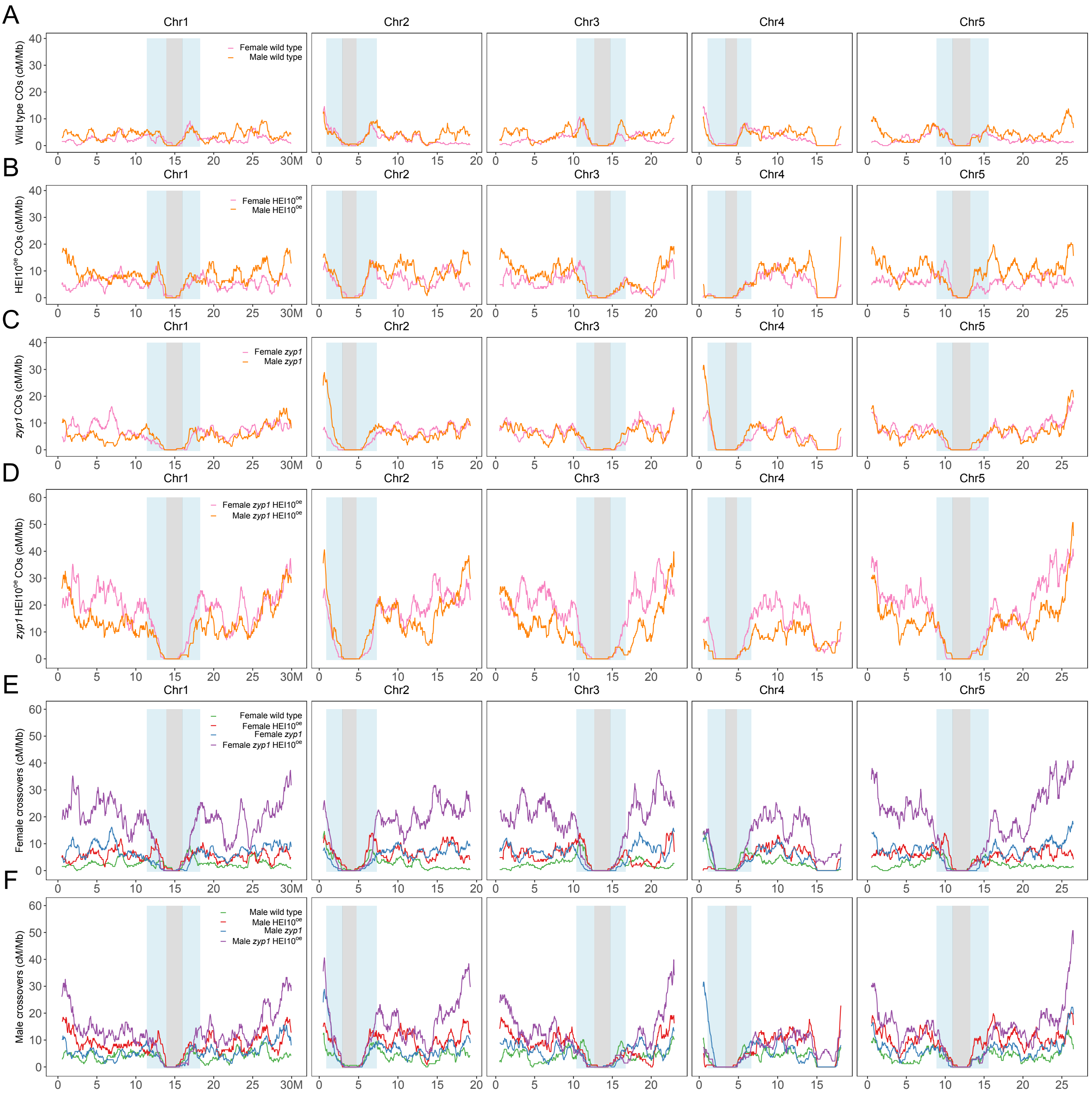


**Figure S3. The distribution of CO frequency along chromosomes.**

Comparison of CO distributions (sliding window-based, with window size of 1 Mb and step size of 50 kb) between female and male of wild type (A), HEI10$^{oe}$ (B), *zyp1* (C) and *zyp1* HEI10$^{oe}$ (D). Comparison of CO distribution among populations in female (E) and male (F) meiosis. The pericentromeric and centromeric regions are indicated by grey and blue shading, respectively. Consistent with previous observations[11,52], there is a ~2.2 Mb region on the long arm of chromosome 4 where recombination is suppressed, which suggests a structural arrangement between the Col and L*er* strains.

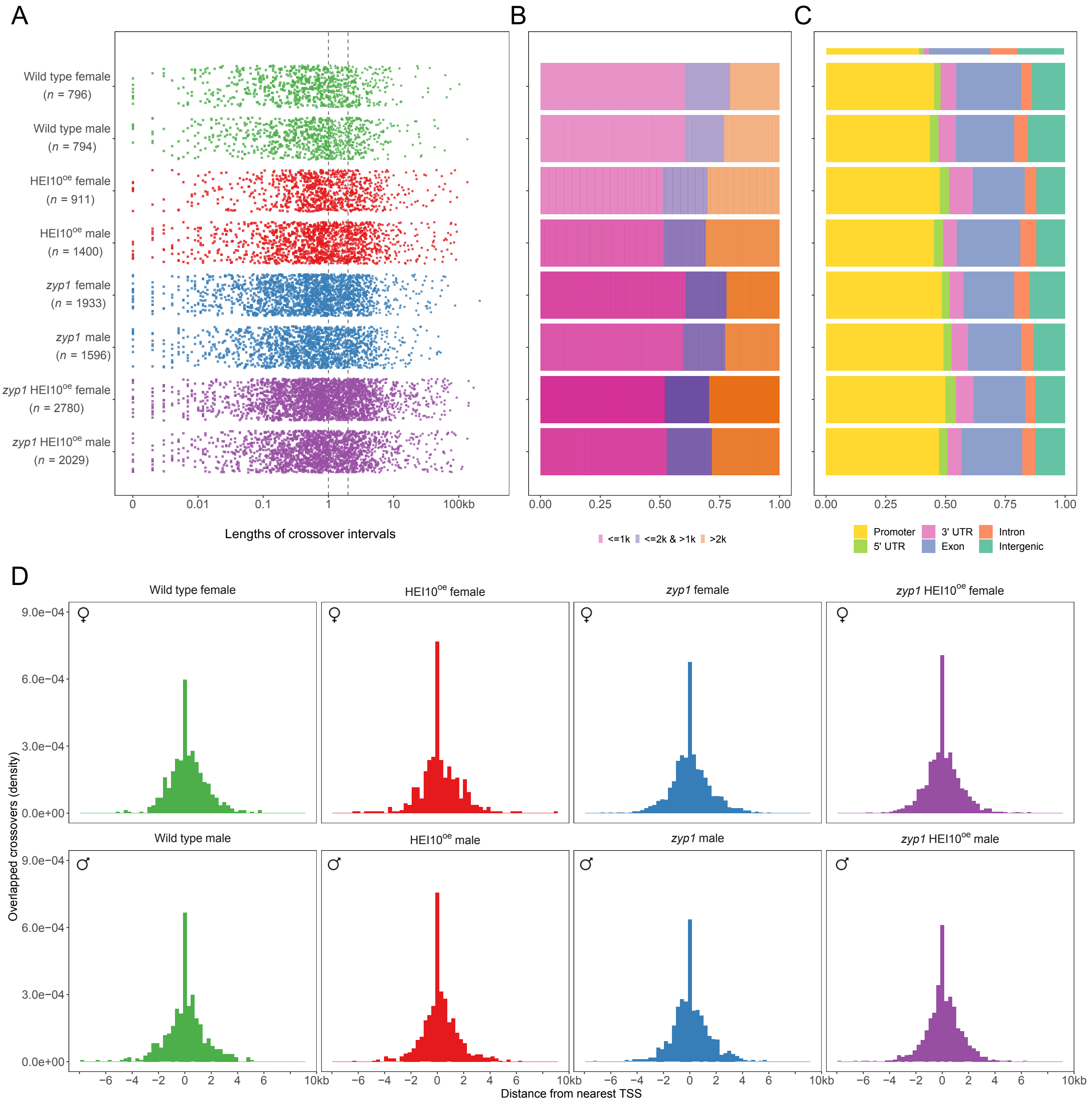


**Figure S4. The fine scale distribution of COs.**
(A) The distribution of the length of CO intervals across populations. The length of 1kb and 2kb are indicated by grey dash lines. The number of analysed COs is shown in parentheses. The median of CO intervals is 819 bp (B) The distribution of proportion of COs with interval length less than 1kb (high-resolution), more than 2kb, and the rest separately. (C) The distribution of proportion of high-resolution COs overlapped with genomic features. The promoter region is defined as the 2kb upstream of the transcription start site. The proportion of the different genomic features is shown as the bar on the top, which defined by following the priority of promoter, 5' UTR, 3' UTR, exon, intron and intergenic regions. (D) The distribution of distance of high-resolution COs from nearest TSS.

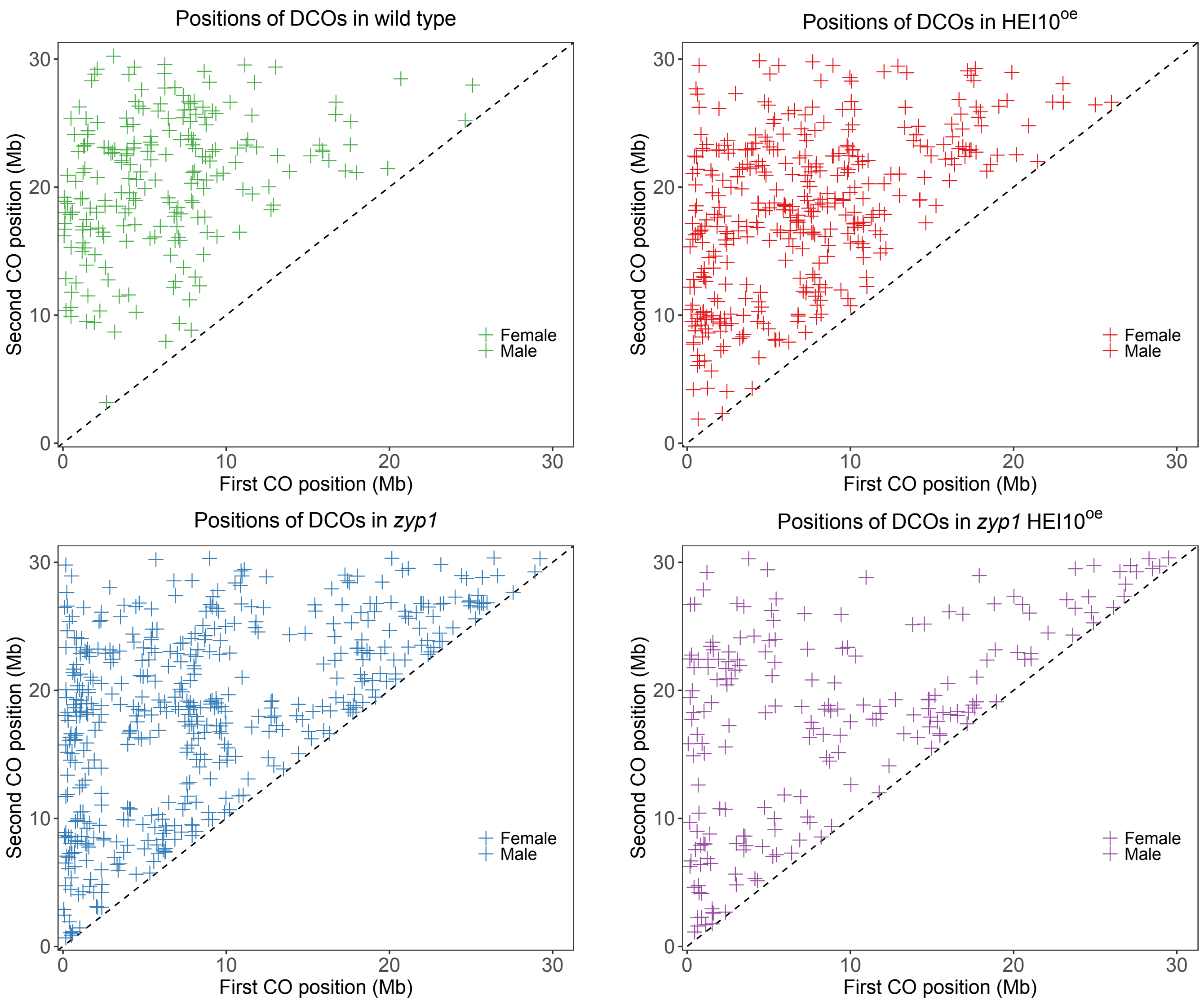


**Figure S5. The distribution of positions of double-COs.**

Relative position of COs for chromosomes with exactly two COs. The position of the first and second CO of the pair, in female and male meiosis of wild type, HEI10$^{oe}$, *zyp1*, and *zyp1* HEI10$^{oe}$, respectively.

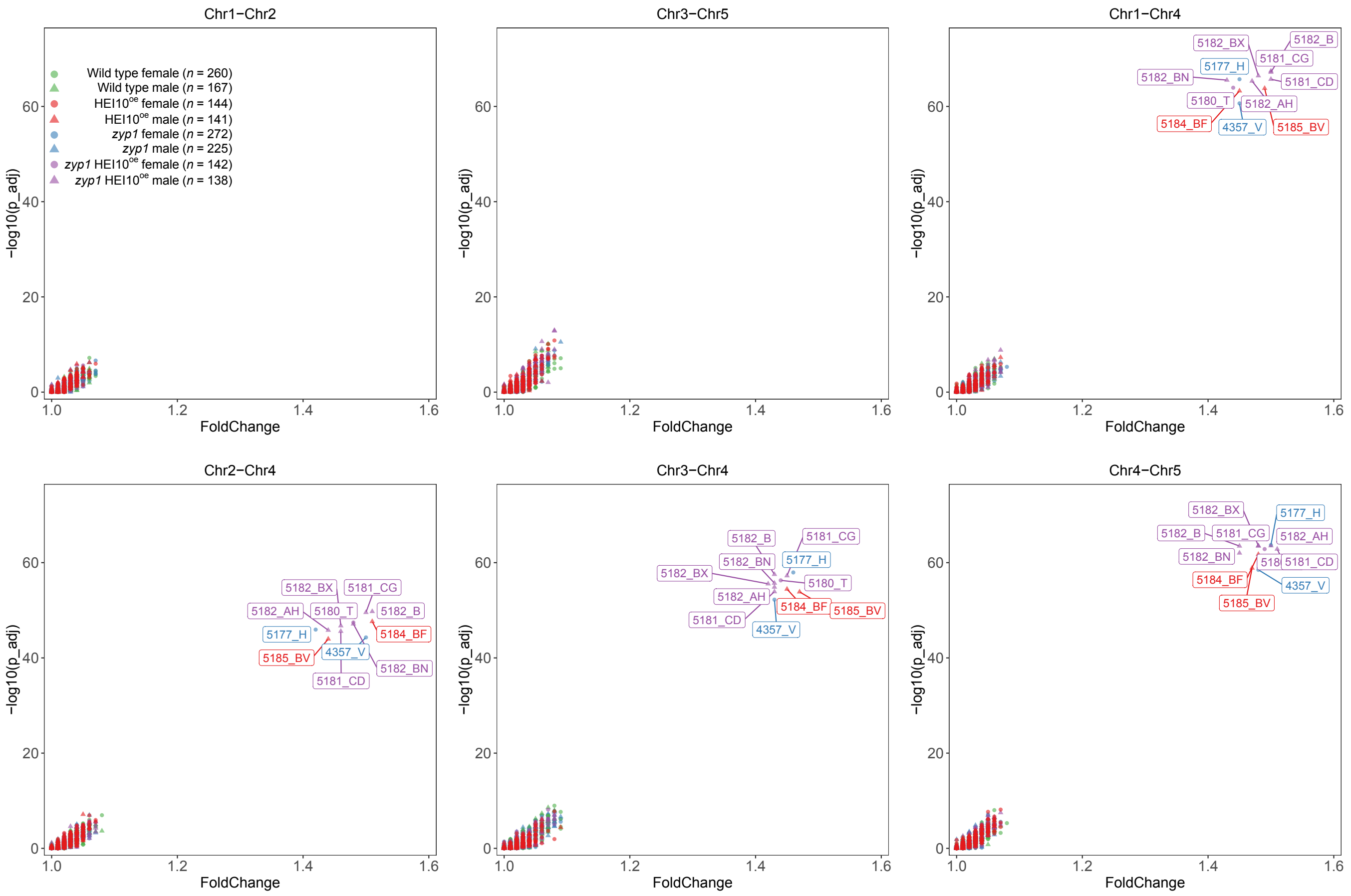


**Figure S6. The analysis of sequencing depths along chromosomes for screening aneuploidy.**
The sequencing depth was calculated for each 100kb non-overlapped interval along chromosomes. The Mann-Whitney test was used for checking the difference between pair of chromosomes, the significant p value was then adjusted by fdr method. The name of the detected aneuploidy is shown with the same color-codes as for corresponding populations.

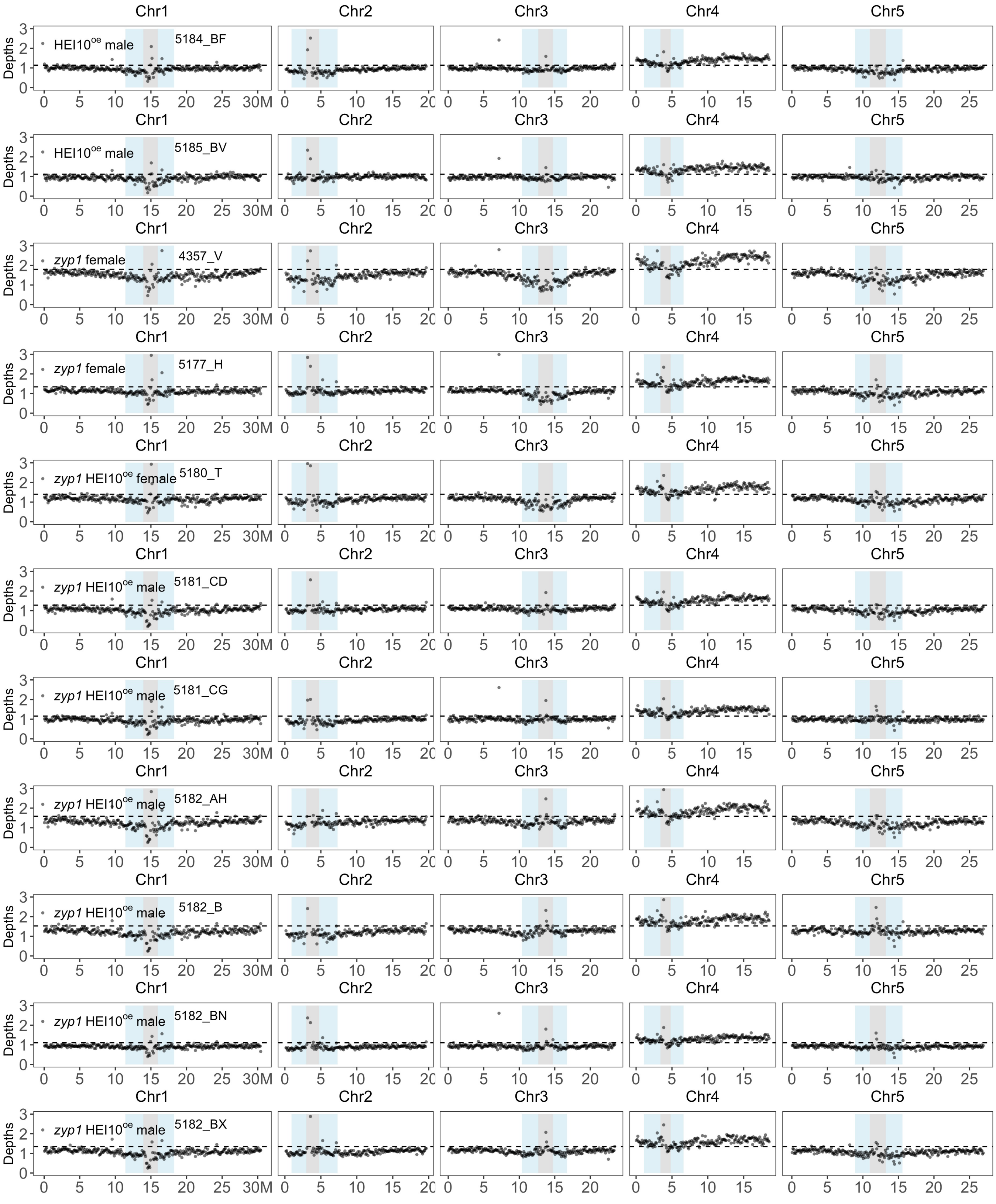


**Figure S7. Sequencing depth along aneuploid chromosomes.**

The sequencing depth was calculated for each 100 kb non-overlapped interval along chromosomes. The pericentromeric and centromeric regions are indicated by grey and blue shading, respectively. The horizontal dashed line indicates the mean sequencing depth of the sample. Aneuploidy is visible by higher coverage of one chromosome compared to the others. The label of the detected aneuploidy and corresponding populations are presented individually.
